# Non-Transgenic Functional Rescue of Neuropeptides

**DOI:** 10.1101/2021.05.10.443513

**Authors:** Elizabeth M. DiLoreto, Douglas K. Reilly, Jagan Srinivasan

## Abstract

Animals constantly respond to changes in their environment and internal states via neuromodulation. Neuropeptide genes modulate neural circuits by encoding either multiple copies of the same neuropeptide or different neuropeptides. This architectural complexity makes it difficult to determine the function of discrete and active neuropeptides. Here, we present a novel genetic tool that facilitates functional analysis of individual peptides. We engineered *Escherichia coli* bacteria to express active peptides and fed loss-of-function *Caenorhabditis elegans* to rescue gene activity. Using this approach, we rescued the activity of different neuropeptide genes with varying lengths and functions: *trh-1, ins-6*, and *pdf-1*. While some peptides are functionally redundant, others exhibited unique and previously uncharacterized functions. The mechanism of peptide uptake is reminiscent of RNA interference, suggesting convergent mechanisms of gene regulation in organisms. Our rescue-by-feeding paradigm provides a high-throughput screening strategy to elucidate the functional landscape of neuropeptide genes regulating different behavioral and physiological processes.

## Introduction

### Neuropeptide Signaling is Complex

When presented with a host of environmental cues, organisms’ sense, interpret, and enact appropriate responses to stimuli ^1,2^. The interpretation and integration of stimuli is a dynamic process that requires neural circuits to be flexible to elicit the proper behavior. For example, a single stimulus can drive multiple reactions depending on internal or environmental states of the organism ^3^. In the roundworm *Caenorhabditis elegans*, when presented with the same small molecular cue (ascr#3), the sexes respond differently: males are attracted while hermaphrodites avoid the cue ^4–6^. Indole elicits different reactions depending on the amount present in the environment. At low concentrations, indole is attractive with a pleasant, floral aroma, though at high concentrations, it is repulsive, smelling pungent, reminiscent of feces or rot ^7,8^.

To modulate competing behaviors in response to the same cue, for example, different neurons can be involved at the circuit level or changes can occur at the synaptic level. A more dynamic approach for modulating neural responses is peptidergic signaling. Neuropeptides are short amino acid chains that serve to regulate neural circuits, while also functioning as neurotransmitters and neurohormones ^9,10^. Neuropeptides signal on longer time scales than, though often in concert with neurotransmitters to regulate synaptic activity, typically through G-protein coupled receptor signaling cascades ^10,11^. Multiple modulators allow neurons to serve unique functions within discrete neural circuits ^12^.

Neuropeptides serve a broad array of functions across the animal kingdom. Oxytocin-like and vasopressin-like peptides are neurohormones that regulate social attachment, lactation, and blood pressure by contracting muscles in mammals, dating back 600-700 million years ^13–18^. However, in the Echinoderm sea star, *Asterias rubens*, vasopressin-like and oxytocin-like neuropeptide ortholog asterotocin serves instead to in relax muscles in the cardiac stomach during fictive feeding ^18^. The *C. elegans* vasopressin ortholog, nematocin, interacts with serotonin and dopamine signaling to modulate gustatory associative memory and male mating behaviors ^13,19^.

*C. elegans* are a microscopic nematode that displays robust behaviors driven by just over 300 neurons ^20,21^. The *C. elegans* genome encodes three classes of neuropeptides: FMRFamide-like peptides (FLP), insulin-like peptides (INS), and non-FLP/insulin neuropeptide-like peptides (NLP) ^22–24^, encoding over 300 individual neuropeptides through 131 genes that modulate the functional connectome ^9,25,26^. The complexity of the neuropeptide genome, combined with extra-synaptic neuropeptides signaling makes elucidating the role of individual peptides difficult, as canonical studies often rely on null mutations and transgenic rescues ^27–30^. As such, these studies are often incomplete, as full gene rescue restores complete preproproteins and makes discrimination of discrete peptide function impossible ^9,31^. Here we present a strategy that rescues neuropeptides synthesized endogenously within the worm genome via engineered bacteria expressing individual peptides.

### Development of Rescue-by-Feeding Paradigm

Current methods of genetic rescue in *Caenorhabditis elegans* are not ideal for understanding the function of individual peptides; whether by cost inhibition or equipment limitations ^32,33^ or due to whole gene rescue not being ideal for studying discrete peptides ^34–37^ Feeding of peptides via *E. coli* has proven successful in manipulating *C. elegans* biological function. Feeding the scorpion venom protein, mBmKTX, modulates lifespan and egg laying behaviors ^38^.

Neuropeptide rescue-by feeding, outlined here, expands on RNA interference (RNAi) feeding paradigms, which have been successfully used to test gene function ^39,40^, wherein a plasmid encoding the RNA of interest is driven by isopropyl-β-D-thiogalactoside (IPTG) −induction. Wherein RNAi employs paired T7 promoters facing one another to produce double-stranded RNA, our protocol uses one T7 promoter to generate mRNA encoding the peptide of interest. The use of Gateway Cloning to develop expression vectors allows for the development of high-throughput experiments targeting individual peptides, or combinations thereof ^41,42^ (**Figure 1a**). Additionally, this technique is readily accessible compared to traditional transgenic rescue approaches.

**Figure 1.**
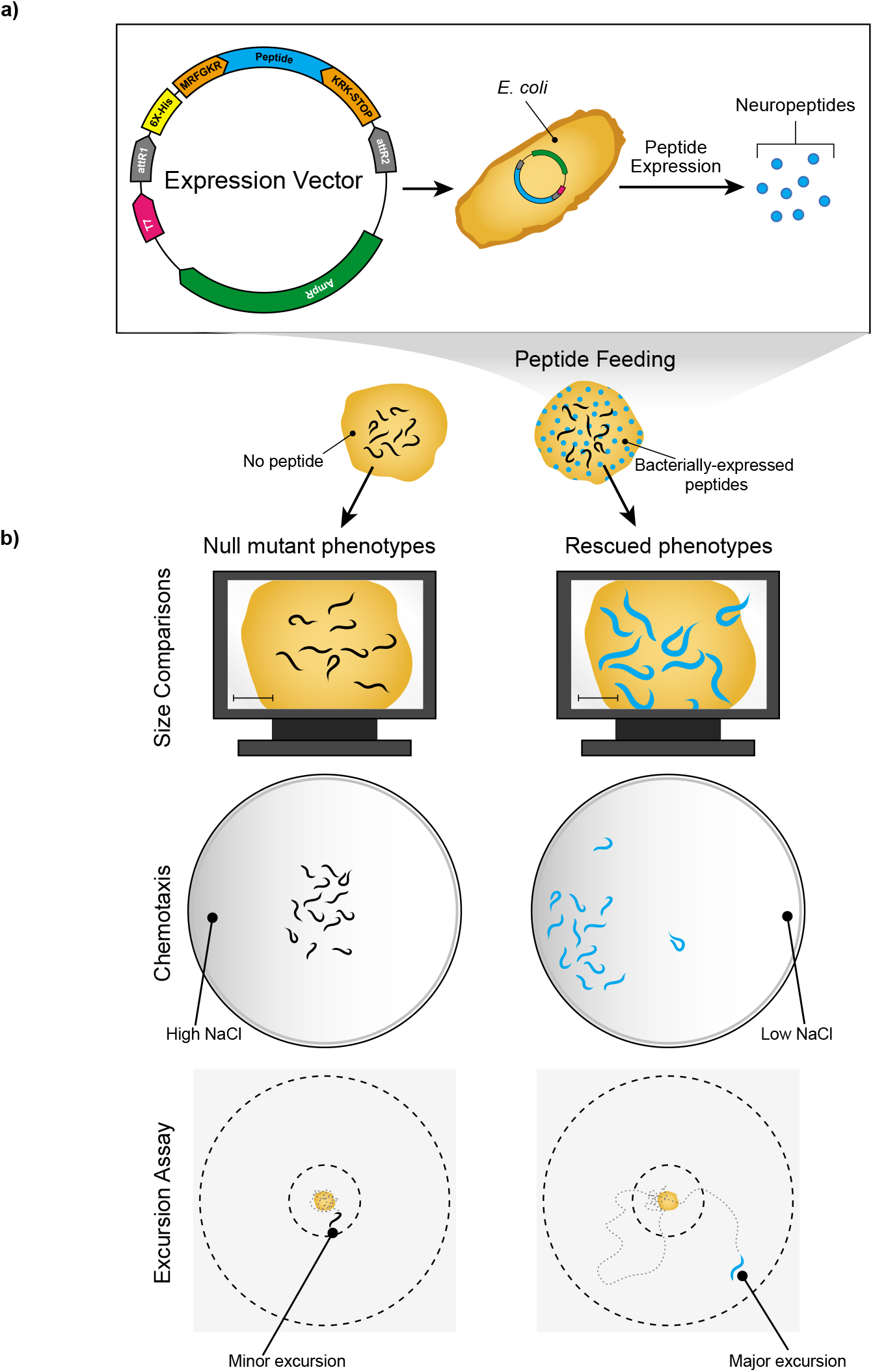
Overview of the Rescue-By-Feeding Technique to Rescue Neuropeptide Function. **a)** Design of an expression vector containing the peptide sequence of interest. The vector sequence includes Gateway Cloning sites attR1 and attR2, a 6x-histidine tag (6x-His), and MRFGKR and KRK-STOP sequences flanking the peptide sequence of interest. This vector is transformed into *E. coli* DH5α cells allowing for the expression of peptides. The bacteria expressing the neuropeptides is then fed to neuropeptide loss-of-function worms and assayed for rescue of behavior. **b)** Different assays used to quantify behavioral activity. We used three different assays for the quantification of rescue; (*1*) Size Comparison, (*2*) Chemotaxis and (*3*) Excursion Assay. For Size Comparison, mutant worms are fed the peptide for 48 hours before assaying for body morphology on an automated worm tracker. In the second assay, Chemotaxis, we used an NaCl gradient chemotaxis assay to salt attraction of worms of rescued neuropeptides. The Excursion Assay quantified the ability for males to leave a food source in search of mates. Minor Excursion denotes worms that did not leave the food source. Major Excursion denotes worms that left the food source in search of mates, indicating rescue (See **<u>Methods</u>** for details of the assays).

The paradigm rescues individual, processed neuropeptides, leveraging the genetic amenability of the *C. elegans* food source, *Escherichia coli*, to circumvent the need for transgenic development and enable high-throughput rescue and elucidation of individual neuropeptide function. We demonstrate the application of this paradigm by rescuing behaviors driven by neuropeptides synthesized from *trh-1, ins-6*, and *pdf-1* (**Figure 1b**).

## Results and Discussion

### Functional Redundancy of Thyrotropin-Releasing Hormone (TRH)-like Peptides in *C. elegans*

Recent work has revealed that *C. elegans* express homologs of mammalian thyrotropin-releasing hormone (TRH) ^43^. Expression of the nematode gene, *trh-1*, in the pharyngeal motor neurons M4 and M5 results in the production of a TRH-like neuropeptide precursor that is processed into two matured peptides: TRH-1A (GRELF-NH_2_), and TRH-1B (ANELF-NH_2_) ^31,43^. Like FLP and other NLP peptides, these peptides are also flanked by di/tribasic residues, allowing them to be processed by EGL-3 ^44,45^.

In mammals, TRH is essential for proper metabolism and growth ^46^, and can even induce metamorphosis in certain amphibians ^47^. The initial characterization of *trh-1* and the cognate receptor gene, *trhr-1*, revealed a similar role within the *C. elegans* nervous system ^43^. Animals expressing a *trh-1* pro-peptide that is truncated prior to peptide translation due to an 8-bp indel-frameshift are significantly shorter and thinner than their wild-type counterparts, resulting in a relative body volume defect (**Figure 1b**, Size Comparisons), 48-hours after larval stage 1 (L1) arrest (Mann-Whitney test, *p* = 0.0012) (**Figure 2a**) ^43^. *In-vitro* studies involving biochemical activation of the TRHR-1 receptor by either peptide (TRH-1A or TRH-1B) suggests functional redundancy of these peptides in rescuing body volume defect in *trh-1* animals ^43^.

**Figure 2.**
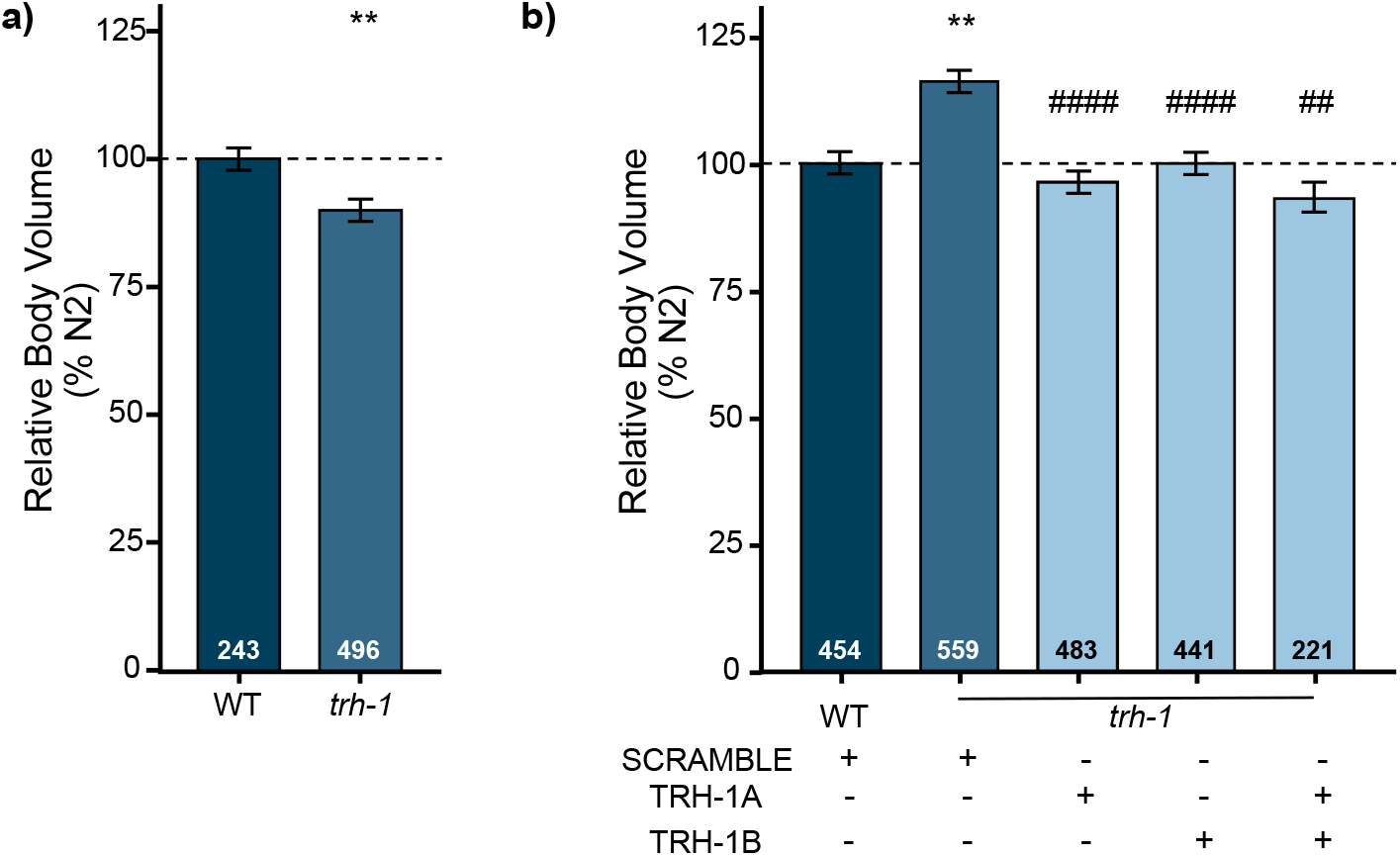
Feeding of TRH-1 Peptides Rescues the Body Volume Defects in *trh-1* Mutants. **a)** *trh-1* mutant worms exhibit body volume defects. *trh-1* mutants show a reduced body volume compared with wild-type worms when fed *E. coli* OP50 (Mann-Whitney test, ** *p* = 0.0012). **b)** Feeding of TRH-1 peptides results in restoration of body volume in *trh-1 lof* worms. *trh-1* mutant worms fed with scramble peptide, display significant increase in body volume feeding compared to wild-type animals. *trh-1* mutants fed either TRH-1A or TRH-1B or even a combination of both exhibit a restoration of wild-type body volume. Error bars denote SEM. *n* values denoted in graph. Kruskal-Wallis followed by Dunn’s multiple comparisons, **/## *p* < 0.01, #### *p* < 0.0001, *denote mutant worms compared to wild-type control, #denote mutant worm condition compared to mutant worms fed scramble peptide.

We cloned both TRH-1 peptides; TRH-1A and TRH-1B in our expression vector as described in **Figure 1**. *trh-1 loss-of-function* (*lof*) mutant animals fed scramble peptide (SCRAMBLE, NSKLHRGGGRSRTSGSTGSMASHARGSPGLQ-NH2) expressed in *E. coli* DH5α cells experience a significant increase in their body volume compared to wild-type animals (Kruskal-Wallis followed by Dunn’s multiple comparison test, *p* = 0.0015) (**Figure 2b**). However, feeding *trh-1 lof* animals with *E. coli* bacteria expressing either TRH-1A, or TRH-1B (*trh-1 lof* versus TRH-1A or TRH-1B fed, *p* < 0.0001) or a combination of both TRH-1A and TRH-1B peptides resulted in a complete restoration of wild-type body volume (*trh-1 lof* versus TRH-1A+B fed, *p* = 0.0019; wild-type versus TRH-1A+B fed, *p* > 0.9999) (**Figure 2b, Supp. Figure 1a-b, c-h** for length, width and area). This suggests that feeding bacteria expressing TRH-1A or TRH-1B, or a combination of the two peptides, to *trh-1* mutant animals results in rescue of body volume, suggesting that they are functionally redundant. While transgenic *C. elegans* expressing the fulllength *trh-1* gene under its endogenous promoter restores wild-type body volume ^43^, the role of individual peptides (TRH-1A and TRH-1B) in regulating this phenotype was not addressed. Our rescue-by-feeding strategy offers an easy, high-throughput *in vivo* biological confirmation of function.

An interesting observation we noticed is that the quality of food determines the body morphology phenotype in *trh-1* mutant worms (**Figure 2a, b**). *trh-1* mutant worms when fed on the standard food source, *E.coli* OP50, displayed reduced body volume as previously published (**Figure 2a**) ^43^. However, *trh-1 lof* worms reared on *E. coli* DH5α cells exhibited increased body volume compared to wildtype animals raised under similar conditions (**Figure 2b**). In addition, we observed that other body morphology characteristics such as relative body length (**Supp. Figure 1c, d**), width (**Supp. Figure 1e, f**), and area (**Supp. Figure 1g, h**) are different in worms fed with *E. coli* DH5α compared to *E. coli* OP50 fed animals. *E. coli* OP50 strain, an uracil auxotroph is the most commonly used laboratory bacterial food source as it grows in thin lawns which allow easier visualization of worms ^48^. However, studies have shown that *C. elegans* prefer other more nutritious bacteria such as HB101 or *Comamonas* sp. for its nutrition ^49^. We propose that the reduced body volume defect in *trh-1 lof* worms reared on *E. coli* OP50 could be a result of an inefficiency of nutrient absorption due to a difference in the nutrient composition of the two *E. coli* strains DH5α and OP50. Our studies corroborate previous literature, wherein *trh-1* mutant worms reared on *E. coli* HB101 exhibit no defect in relative body volume in compared to *E. coli* OP50 fed animals ^43^.

### Feeding of INS-6 Peptides Rescues Chemotaxis Defects Exhibited by *ins-6* Mutants

Insulin and insulin-like peptides serve signaling functions in *Drosophila* and *C. elegans* homologous to the human insulin-like growth factor (IGF), which regulates FOXO activity ^50^. In *C. elegans, ins-6* encodes only one processed INS-6 peptide (VPAPGETRACGRKLISLVMAVCGDLCNPQEGKDIATECCGNQCSDDYIRSACCP-NH2) ^51,52^, though the gene was originally postulated to encode two putative proteins ^9,23^. *ins-6* functions in dauer formation ^51,53^ and sensory modulation of large fluctuations in salt concentration, with loss of *ins-6* causing dysfunction of NaCl attraction (**Figure 1b**, Chemotaxis) ^54^

While wild-type worms were attracted to high concentrations of salt (750 mM), there was a significant decrease in attraction in *ins-6* mutant animals, as measured by a Chemotaxis Index (CI) (Paired *t*-test with samples of equal variance, *p* = 0.0248) (**Figure 3a**) ^54^ Salt attraction was partially rescued with genetic reintroduction of *ins-6* under its endogenous promotor (**Figure 3a**). Similarly, the CI of *ins-6 lof* animals fed scramble peptide was significantly different from the wild-type animals fed scramble (**Figure 3b**). Feeding of *E. coli* expressing INS-6 peptide to *ins-6 lof* animals resulted in a complete restoration of attraction to high salt concentration (ANOVA, followed by Bonferroni’s Correction, *ins-6 lof* fed scramble versus INS-6 fed, *p* = 0.0101) (**Figure 3b**).

**Figure 3.**
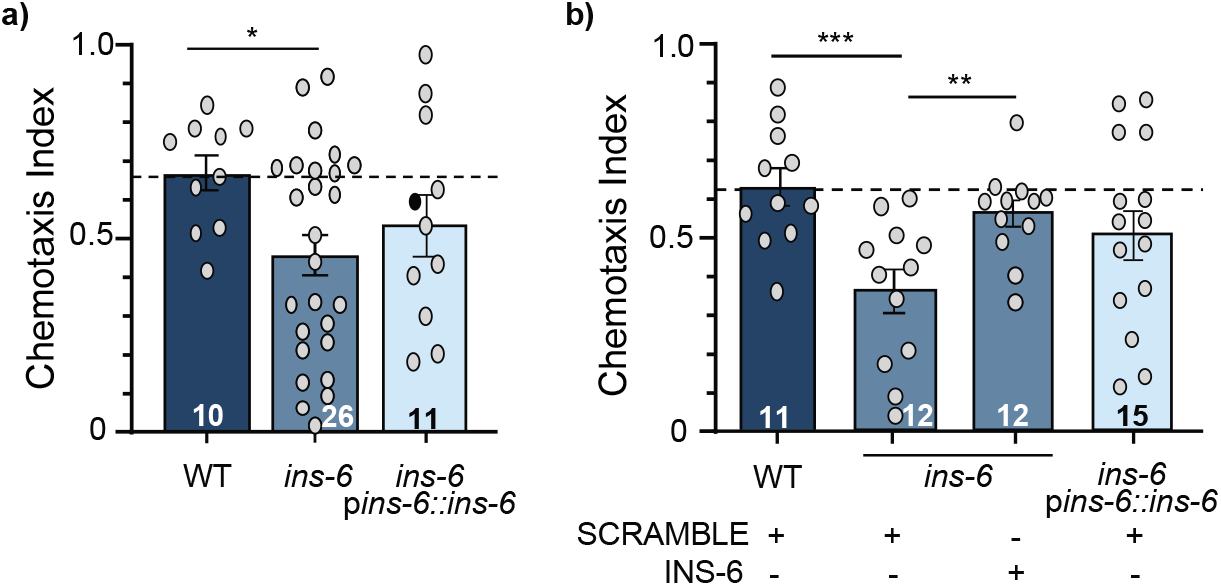
Chemotaxis Defects of *ins-6* Neuropeptide Mutants are Rescued by Feeding the INS-6 Peptide. **a)** Chemotaxis Index to 750 mM NaCl of wild-type (*WT*), *ins-6* loss of function, and genetic *ins-6* rescue (*ins-6;Pins-6;ins-6*), animals fed *E. coli* OP50. Animals with *loss-of-function ins-6* gene exhibit significant decreased attraction and genetic rescue of *ins-6* results in a partial rescue of chemotaxis. **b)** Chemotaxis Index of peptide fed worms to 750 mM NaCl. *ins-6 lof* worms fed scramble display significantly lower chemotaxis compared to scramble-fed wild type animals. Feeding of INS-6 peptide to ins-6 *lof* worms results in complete rescue of NaCl chemotaxis. Overexpression of INS-6 does not result in changes in attraction towards 750 mM NaCl. *n* values are denoted in graphs. Error bars denote SEM. One-Way ANOVA, followed by Bonferroni’s Correction * *p* < 0.05, ** *p* < 0.01, *** *p* < 0.001, **** *p* < 0.0001, comparisons noted with line over bars.

Overexpression of INS-6 did not increase chemotaxis towards high salt, as wild-type animals fed INS-6, along with a full *ins-6* genetic rescue, exhibited slight defects in CI towards high salt (ANOVA, followed by Bonferroni’s Correction, wild-type versus fed, *p* = 0.08071; wild-type versus transgenic overexpression, *p* = 0.2403) (**Supp. Figure 2b**). Partial genetic rescue of *ins-6* limited to the ASI sensory neuron (*ins-6;ASI::ins-6*) was sufficient to rescue neurophysiological function of AWC^ON^ NaCl sensation ^54^, though it was not sufficient for rescuing behavioral attraction to salt (ANOVA, followed by Bonferroni’s Correction, *p* = 0.2136) (**Supp. Figure 2b**). While our rescue-by-feeding paradigm was able to rescue the behavioral phenotypes of aberrant *ins-6* signaling, the ability to combine this method with calcium imaging techniques is still in development.

### Differential Functional Activity of Peptides Encoded by Pigment Dispersing Factor (*pdf*)-1

Wild-type males display a characteristic exploratory behavior when left on a lawn of food, as well-fed males leave food in search of mates ^55,56^. The pigment dispersing factor *pdf-1* neuropeptide plays a significant role in male mate-searching behavior, balancing the neural circuits controlling two predominant male interests: finding food and finding mates ^55,57^. Like *trh-1*, the *pdf-1* precursor encodes two neuropeptides: PDF-1A (SNAELINGLIGMDLGKLSAVG-NH2) and PDF-1B (SNAELINGLLSMNLNKLSGAG-NH2) ^58,59^.

We quantified the behavioral activity of *pdf-1 lof* males using a food-leaving behavior as previously described ^55,60^. Individuals were placed on a small food spot, and track patterns were scored at three different time points (2, 6, 24 hrs) (**Figure 4a**). ‘‘Never left food’’ indicate the absence of tracks outside the food spot. ‘‘Minor excursion’’ indicates that tracks were observed not beyond 1 cm from the food. ‘‘Major excursion’’ indicates the presence of tracks past the 1 cm boundary (**Figure 1b**, Excursion Assay). We additionally quantified this food-leaving behavior as a measure of mate-searching behavior. In this behavior well-fed males left food in search of mates and the data is represented as a Probability of Leaving (P_L_) (**Supp. Figure 3a, b**) ^55,56^.

**Figure 4.**
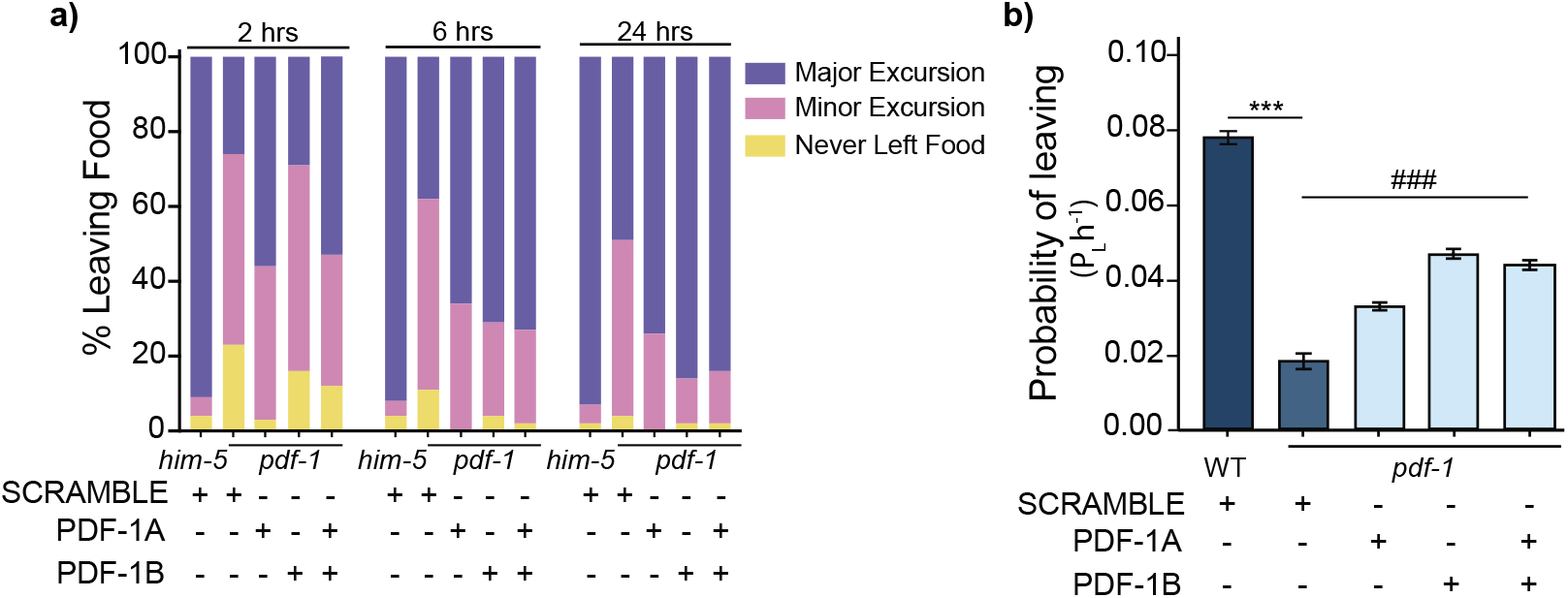
*pdf-1 lof* males fed discrete PDF-1 peptides differentially rescue food-leaving behavior. **a)** Percent of male worms leaving food across time is used to visualize the location of the males a different time points. *him-5* males serve as the *wild-type* worms in this assay compared to males deficient in *pdf-1* fed PDF-1 peptide to rescue behavior. At each time point the percent of worm within each boundary is noted (Major Excursion is beyond 10 mm radius of the food (purple), Minor Excursion is worms who left food but stayed within 10 mm (pink), and those remaining on food were classified as Never Left Food (yellow)). **b)** Probability of Leaving. To compare the leaving behavior of the worms as a likelihood of the worms seeking to travel beyond 35 mm from the food. *n; him-5* males fed scramble = 55, *pdf-1* males fed scramble = 53, *pdf-1* males fed PDF-1A = 58, *pdf-1* males fed PDF-1B = 51, *pdf-1* males fed PDF-1A and PDF-1B = 51). Error bars denote SEM. ANOVA, followed by Bonferroni’s Correction, ***/###*p* < 0.001, #### *p* < 0.0001, *denote mutant worms compared to wild-type control, #denote mutant worm condition compared to mutant worms fed scramble peptide.

*pdf-1 lof* males did not leave food as readily as wild-type worms suggesting that these worms do not display exploratory behavior, as previously described (Paired *t*-test with samples of equal variance, *p* < 0.0001) (**Supp. Figure 3b, c**) ^55^. When fed either PDF-1 peptide (or a 1:1 combination of both), *pdf-1 lof* males had a higher P_L_ compared to scramble fed *pdf-1 lof* males (ANOVA, followed by Bonferroni’s Correction,*p* < 0.0001) (**Figure 4b**). However, PDF-1B and a 1:1 ratio of both peptides resulted in a significantly different P_L_ from *pdf-1* males fed PDF-1A alone (ANOVA, followed by Bonferroni’s Correction, *p* < 0.0001). Rescue-by-feeding resulted in a partial rescue: while the P_L_ of peptide-fed males were significantly higher than *pdf-1 lof* males fed scramble, they were still significantly lower than wild-type males fed scramble peptide (ANOVA, followed by Bonferroni’s Correction, *p* < 0.0001) (**Figure 4b**). Future studies employing this paradigm can now focus on teasing apart the difference in PDF-1A and PDF-1B function in mate-finding behavior, as we have successfully employed in dissecting FLP-3 peptide functions ^61^.

We present here a method of neuropeptide rescue that exploits bacterial expression to deliver individual peptides to rescue behavioral and growth-related phenotypes. We show that while some peptides are redundant in function (**Figure 2**), others rescue phenotypes driven by large peptides (**Figure 3**), while novel functions are displayed by other peptides over long timescales (**Figure 4**). Our novel technology to rescue peptide function is advantageous over transgenic studies, as it allows for functional characterization of individual peptides. This genetic tool is built off the principles of RNAi feeding techniques to supply worms with the peptide of interest through their food source as previously shown for the scorpion venom peptide mBmKTX to alter lifespan and egg-laying behavior in *C. elegans* ^38^. More recently, we have improved upon this paradigm to characterize the FMRFamide-like peptide gene, *flp-3* ^61^. This gene encodes ten different peptides and we leveraged the rescue-by-feeding technology to elucidate that only two of the ten peptides encoded by the precursor are active in controlling the behavioral response of males to a mating pheromone ^61^. These studies support the assertion that feeding peptides to *C. elegans* via their *E. coli* food source is sufficient to rescue mutant phenotypes.

Based on our results, we propose that the neuropeptide rescue-by-feeding strategy delivers mRNA ready for translation by the *C. elegans* cellular machinery, rather than supply *C. elegans* with fully translated and processed peptides. The plasma membrane of a cell is an intricate complex of multiple lipid and protein molecules. Small molecules with moderate polarity diffuse through the cell membrane passively, but most metabolites and short peptides require specialized membrane transporters for translocation ^62^. Given the large number of neuropeptides encoded in the genome of *C. elegans*^9^, having specialized transporters for peptide transport is not feasible ^63^. The strongest piece of evidence for this statement lies in that the processed INS-6 peptide is 54 amino acids in length: the *C. elegans* intestine expresses peptide transporters that only uptake smaller, inactive di- and tri-amino acid peptide chains ^63^. Thus, rescue of chemotaxis behavior by INS-6 peptide feeding (**Figure 3b**) suggests that the mechanism of rescue may be similar to double-stranded RNA (dsRNA) uptake, rather than peptidergic transport across the intestinal membrane ^64,65^. Furthermore, previous studies have shown that dsRNA exhibit transgenerational inheritance ^66^, despite degradation rates,. Our rescue of the exploratory behavior of *pdf-1 lof* males even in the 24-hour timescale (**Figure 4**) suggest that our feeding paradigm exploits similar mechanisms, allowing peptide-encoding mRNAs to remain present throughout the assay. We propose that if peptides were taken up, their degradation rates would likely impede rescue efficiency at later timepoints unlike what we observe in our experiments suggesting that the mechanism is similar to double-stranded RNA (dsRNA) feeding, with mRNA uptake driving rescue rather than peptide uptake.

Feeding *E. coli* DH5α expressing peptides results in altered phenotypes compared to *E. coli* OP50 (**Figure 2, 3**) and (Reilly *et al*., in review) ^61^. Future studies will focus towards optimizing experimental protocols for feeding. Two areas of optimization we envision involve designing an expression vector strategy that can multiplex multiple neuropeptide genes and finding the “best” food source for the rescue experiments. Given the differences in nutrient composition of the different *E. coli* strains and worms preferring more nutritious bacteria ^49^, expressing peptides in HB101 or HB115 strains offer viable alternatives.

Understanding neuropeptide function is essential both from a perspective of regulation of neuronal (synaptic and non-synaptic) channels of communication, and form a global view of general neuronal functional assignments throughout the brain. Revealing how a particular neuropeptide acts both at the cellular and subcellular peptide receptor level is critical link in understanding the role of neuromodulation of circuits. An enhanced experimental pipeline to investigate peptide function will enable progress toward answering how neural circuit activity within the network in its different states and identification are modulated by neuropeptides, resulting in flexible decision-making during behaviors.

## Methods

### *C. elegans* strains

The *C. elegans* strains N2 and CB4088 (*him-5(e1490)*) were obtained from the *Caenorhabditis* Genetics Center. The LSC1118 (*trh-1(lst1118)*) strain was generously provided by Isabel Beets at KU Leuven for the body morphology tests. The strains used in the Chemotaxis Assay were gifted by Prof. Shreekanth Chalasani, Salk Insitutute (IV302 (*ins-6(tm2416)*; kyEx2595) at the Salk Institute, and Yun Zhang (ZC239 (*ins-6(tm2416)II; yxEx175[Pins-6::ins-6; Punc-122::gfp*]) at Harvard University. The UR954 (*pdf-1(tm1886);him-5(e1490)*) strain was provided by Douglas Portman at the University of Rochester.

### Peptide plasmid design and generation

DNA sequences encoding individual peptides were identified via www.wormbase.org. Sequences were flanked with the endogenous cleavage sites for the EGL-3 processing enzyme, which cleaves dibasic resides. Sequences encoding MRFGKR and KRK-STOP codons were therefore placed prior to, and following the peptide codon sequences, respectively. Finally, Gateway Cloning sites attB1 and attB2 sites were attached to the ends of the sequences. These final sequences (comprised of attB1::MRFGKR::peptide::KRK-STOP::attB2) were ordered from IDT (Integrated DNA Technologies) using their DNA Oligo and Ultramer DNA Oligo services, depending on the size of the oligo ordered. Both forward and reverse sequences were ordered (**Supp. Table 1a**).

Lyophilized oligos were prepared following IDT Annealing Oligonucleotides Protocol (https://www.idtdna.com/pages/education/decoded/article/annealing-oligonucleotides). In short, they were resuspended in Duplex Buffer (100 mM Potassium Acetate; 30 mM HEPES, pH 7.5; available from IDT), preheated to 94 °C to a final concentration of 40 μM. Complimentary oligo sequences were then mixed in equimolar ratios, and placed in a thermocycler at 94 °C for 2 minutes, prior to a stepwise cooling to room temperature.

Annealed oligos were used to perform a BP reaction with pDONR p1-2 donor vector to generate pENTRY clones (Gateway Cloning, ThermoFisher). Entry clones were then recombined with pDEST-527 (a gift from Dominic Esposito (Addgene plasmid # 11518)) in LR reactions generating expression clones. The scramble control was generated in an identical manner, with the sequence between the cleavage sites being amplified from pL4440 (provided by Victor Ambros, University of Massachusetts Medical School, MA) (**Supp. Table 1b**). Purified plasmids were stored at −20 °C or −80 °C for long term storage.

### Peptide Plate Preparation

The expression clone for both the peptide of interest and the control was transformed into competent DH5α cells (NEB^®^ [New England Biolabs] 5-alpha Competent *E. coli* (High Efficiency) Cat# C2987I).

Peptide-expressing cultures on LB agar plates (10 g/L NaCl, 10 g/L Tryptone, 5 g/L Yeast Extract, 5 g/L Agar) containing 100 μg/μL ampicillin (AMP), were stored at 4 °C for up to 2 months. From this plate, single colonies were selected for growth in 5 mL of LB media containing ampicillin at 37 °C for 16 hours.

The optical density of the grown cultures (OD_600_) was adjusted to 1.0. 75 μL of 1.0 OD_600_ peptide-expressing culture was plated onto 60 mm NGM agar plates (50 mM NaCl. 25 mM KPO_4_ (pH 6.0), 1mM MgSO_4_, 1mM CaCl_2_, 5 μg/L cholesterol, 17 g/L agar, 2.5 g/L peptone) containing 100 μg/μL AMP and 1 mM isopropyl-β-D-thiogalactoside (IPTG) at room temperature for at least 8 hours prior to use, for no longer than 1 week.

### Peptide Feeding

All animals were maintained on bacterial lawns of OP50 *E. coli*, at 20 °C, on NGM agar plates until the start of experiments. Mutant or control animals were transferred onto lawns of DH5α *E. coli* expressing either scramble peptide or the rescue peptide-of-interest on NGM plates containing 100 μg/μL AMP and 1 mM IPTG. Animals were reared on the peptide-expressing lawns for at least 48 hours at 20 °C before assaying for rescue of mutant phenotype at the appropriate developmental stage. See each section of the behavioral assay methods for the exact peptide feeding parameters.

### Size Comparison (*trh-1*)

The body morphology different of *trh-1* mutants was compared to wild-type worms by body volume measurements. Wild-type (N2) and *trh-1* animals were synchronized by embryonic starvation following alkaline hypochlorite protocols ^67^. Ten gravid hermaphrodites from OP50 *E. coli* plates were picked into 30 μL drops of alkaline hypochlorite solution (20% sodium hypochlorite, 500 mM KOH). Two drops were placed on either side of an unseeded NGM plate, resulting in 20 animals per plate. Plates were then incubated for 24 hours at 20 °C before L1 larval animals washed with 1 mL M9 (22 mM KH_2_PO_4_, 42.25 mM Na_2_HPO_4_, 85.5 mM NaCl, 1 mM MgSO_4_) per unseeded NGM plate. Allow L1 worms to settle into pellet before removing M9 supernatant. Repeat washing step 2 more times. Plate worm pellet onto NGM with 100 μg/μL AMP and 1 mM IPTG plates containing 75 μL peptide lawns.

After at 48 hours at 20 °C ^43^, animals were recorded using a Basler acA2500-14uM camera using WormLab software (WormLab 4.1; MBF Bioscience, 2017) for 65 seconds. Worm *length, width*, and *area* were calculated within the WormLab software (**Supp. Figure 1 c-h**). As the WormLab worm width metric is an average of cross-sections throughout the length of each worm, this was used to calculate the volume of a worm as a cylinder, with the *Volume* = π * (0.5 * *width*)^2^ * *length* (**Figure 2a, b, Supp. Figure 1a, b**).

### Chemotaxis Assay (*ins-6*)

A salt chemotaxis assay was performed as previously described in ^68,69^, to test the functionality of *ins-6* after rescue-by-feeding. Wild-type (N2) and *ins-6* worms were fed either scramble or INS-6 peptide passing 4-6 L4 larval onto an NGM plate containing 100 μg/μL AMP and 1 mM IPTG with 75 μL of peptide OD_600_ = 1.0. These worms were grown at 20 °C for about 4 days until the progeny of the initial L4 worms were young adults.

The day prior to testing, a thin, 10 mL layer of agar was added to 10 cm petri dishes to generate a 10 cm Chemotaxis Plates (5 mM KPO_4_ (pH 6.0), 1 μM MgSO_4_, 1 μM CaCl_2_, 20 g/L agar, and 8 g/L Difco Nutrient Broth). To prepare the “high salt” plates, 750 mM NaCl was added prior to plate pouring. The high salt plate were stored at 4 °C for up to one month, while the Chemotaxis Plates were made one day prior to testing. To create the high and low salt plugs, the back end of a Pasteur pipette was used to punch a 5 mm plug out of each plate. These 2 plugs were placed on opposite edges of the plate. A 10 mm radius was be drawn around the location of the plugs (**Figure 1** for schematic). Plates were stored, lightly covered, overnight at room temperature (~20 °C, <40% humidity).

The day of the assay, young adult worms, either control worms fed OP50 *E. coli* or peptide-fed worms, were washed off peptide plates with M9. Worms were allowed to settle by gravity for 4 minutes before removing the supernatant, washing the worms. Worms were then washed with Chemotaxis Buffer (5 mM KPO_4_ (pH 6.0), 1 mM MgSO_4_, 1 mM CaCl_2_) three times.

To prepare the Chemotaxis plates for the assay, high and low salt plugs were removed with forceps, and the plates spotted with 1 μL of 0.5 M sodium azide (NaN_3_). Approximately 30 μL of worms in Chemotaxis Buffer were spotted onto the edge of the agar, between the location of the high and low salt gradient (**Supp. Figure 2**). Worms were left to chemotax for 1 hour before scoring the number of worms within the 10 mm radii of high and low salt areas.

### Excursion Assay (*pdf-1*)

The mate searching behavior of *pdf-1* mutants was quantified in a food leaving assay. *him-5* animals were used as wild-type control to ensure the presence of males. *him-5* and *pdf-1* animals were fed OP50, scramble peptide, PDF-1A, PDF-1B, or a 1:1 mixture of PDF-1A and PDF-1B cultures. Four to six larval L4 hermaphrodites were passed onto an NGM plate containing 100 μg/μL AMP and 1 mM IPTG with 75 μL of peptide. These worms were grown at 20 °C for about 4 days until the male progeny of the initial L4 worms were young adults. The day prior to the assay, young adult male *pdf-1* or *him-5* worms were singled onto 60 mm plates containing lawns of either OP50 (NGM plates) or appropriate peptide cultures (NGM containing 100 μg/μL AMP and 1mM IPTG).

To determine the mate searching effects of *pdf-1*, we performed a food-leaving assay as described previously ^55,56,60^, with modifications. In this assay, individuals were placed on a small food spot, and track patterns were scored at the indicated times. Assay plates were prepared one day prior to assaying on 100 mm plates with 10 mL of Leaving Assay media (25 mM KPO_4_ (pH 6.0), 1 mM MgSO_4_, 1mM CaCl_2_, 50 mM NaCl, 2.5 g/L Peptone, 17 g/L Agar). On center of plate, 7 μL of OP50 was added. Plates were lightly covered and stored at room temperature overnight (~20°C, <40% humidity).

On the day of the assay, a single male was plated on the assay plate and placed in a dark, 20°C incubator. At 2, 6, and 24 hours after the worms are plated, plates were scored for male worm tracks. Scored were separated into binned ranges ^60^. “Never left food” indicates the absence of tracks outside the food spot. “Minor excursion” indicates that tracks were observed not beyond 10 mm from the food. “Major excursion” indicates the presence of tracks past the 1 cm boundary. We quantified the food leaving behavior at three different time points (2, 6, and 24 hours) ^55^. (**Figure 4a, Supp. Figure 3b**) ^55^.

We additionally quantified the mate searching defect of *pdf-1* as a Probability of Leaving per hour (P_L_) (**Figure 4b, Supp Figure. 3a, c)** ^56^. In this method, we reduce the analysis of the leaving behavior to a single value, by scoring the worms leaving the food lawn as Leavers (traveling beyond 35 mm from the food source) or Non-Leavers (did not travel further than 35 mm from center of food).

## Statistics and Reproducibility

### Size Comparisons

Two to four plates were recorded per day for at least three days to generate data sets. Control plates of N2 and *trh-1* animals raised on the scramble peptide were grown alongside every rescue feeding. Each “worm track” was considered an *n* = 1. Tracks were excluded if (1) worm tracks less than 10 seconds, or (*2*) worms were further than two standard deviations from the mean metric. Following tests for normality, comparisons were made using either Mann-Whitney tests for direct comparisons between N2 and *trh-1* animals reared on OP50, or Kruskal-Wallis non-parametric ANOVAs and Dunn’s multiple comparisons tests for animals raised on peptide feeding strains. Comparisons were made for metrics: *length* (**Supp. Figure 1c, d**), *width* (**Supp. Figure 1e, f**), *area* (**Supp. Figure 1g, h)**, and *volume* (**Figure 2a, b, and Supp. Figure 1a, b**).

### Chemotaxis Assay

Young adult hermaphrodites are tested in the Chemotaxis Assay with no more than three replicates per day, over a minimum of three days, with 100-200 worms/assay plate. The Chemotaxis Index (CI) was calculated as 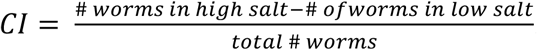. At least 10 plates were assayed in each condition: OP50 control, scramble fed, and INS-6 fed. Plates were excluded from final calculations if (*1*) the total number of worms on the assay plate was <20, or (*2*) if the plate CI was greater than two standard deviations from the average CI. Conditions were compared using Two-tailed *t*-tests with samples of equal variable to compare between two conditions with small samples or ANOVA followed by Bonferroni’s multiple comparison for multiple condition comparisons.

### Excursion Assay

The assay was performed across at least two days per condition, with at least *n* = 50, wherein each assay plate with one male is equal to *n* = 1. In all experiments, the investigators were blinded to genotype. Worm in the >35 mm from the center of food bin were scored as “leavers”^56^. The probability of leaving per hour (Probability of Leaving (P_L_)) was calculated using an R script developed in ^55^, first cited in ^56^, with assistance by (Personal Communications to Barrios, 2020). P_L_ was calculated by the “hazard obtained by fitting an exponential parametric survival model to the censored data using maximum likelihood” (**Supp. Figure 3a**) ^56^. The P_L_h^-1^ values for all *pdf-1* animals fed scramble or peptide rescue were compared to *him-5* worms fed scramble using an ANOVA followed by Bonferroni’s multiple comparison (**Figure 4b, Supp. Figure 3c**).

## Data Availability

The raw data for all figures is available with this manuscript. Additionally, a summary containing the average, n, standard error measure, and associated statistical test is also available.

## Acknowledgements

We would like to thank Jeffrey Marsh for assistance in running the Chemotaxis Assays. We would like to thank Dr. Arantza Barrios for help in executing the leaving assay analysis in R. We extend our thanks to Dr. Shreekanth Chalasani and Dr. Douglas Portman for their insights and updates to method for testing the phenotypes related to *ins-6* and *pdf-1*, respectively. Thank you to Dr. Isabel Beets for her insights into the *trh-1* analysis. This work was funded by NIH R01DC016058 (JS).

## Author’s Contribution

EMD, DKR and JS designed all experiments, EMD and DKR performed all experiments and statistical analysis. EMD co-wrote the paper with DKR and JS.

## Ethics Declaration

The authors declare no competing financial or non-financial interests as defined by Nature Research.

---

**Supplementary Figure - 1.**
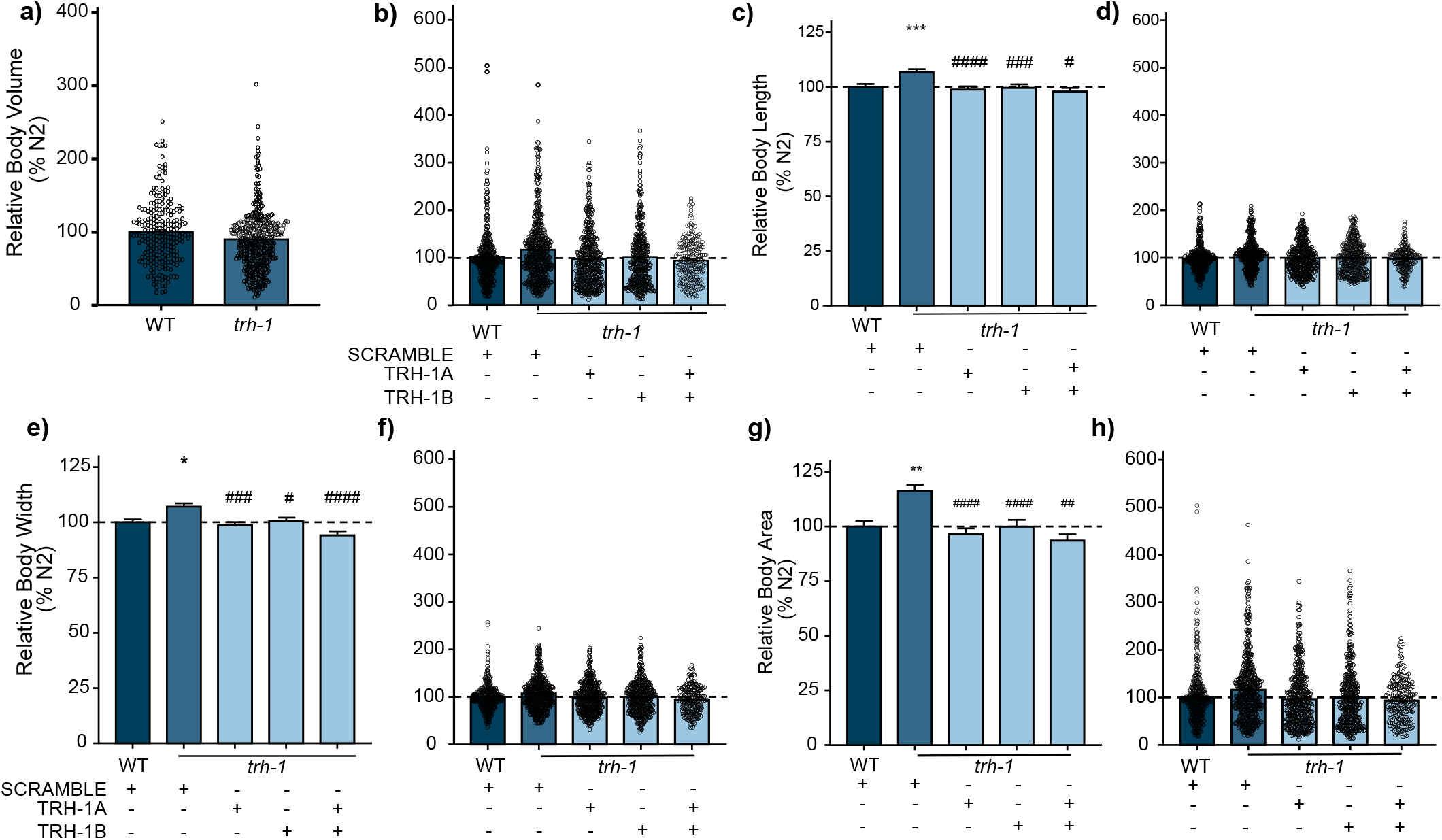

**Supplementary Figure - 2.**
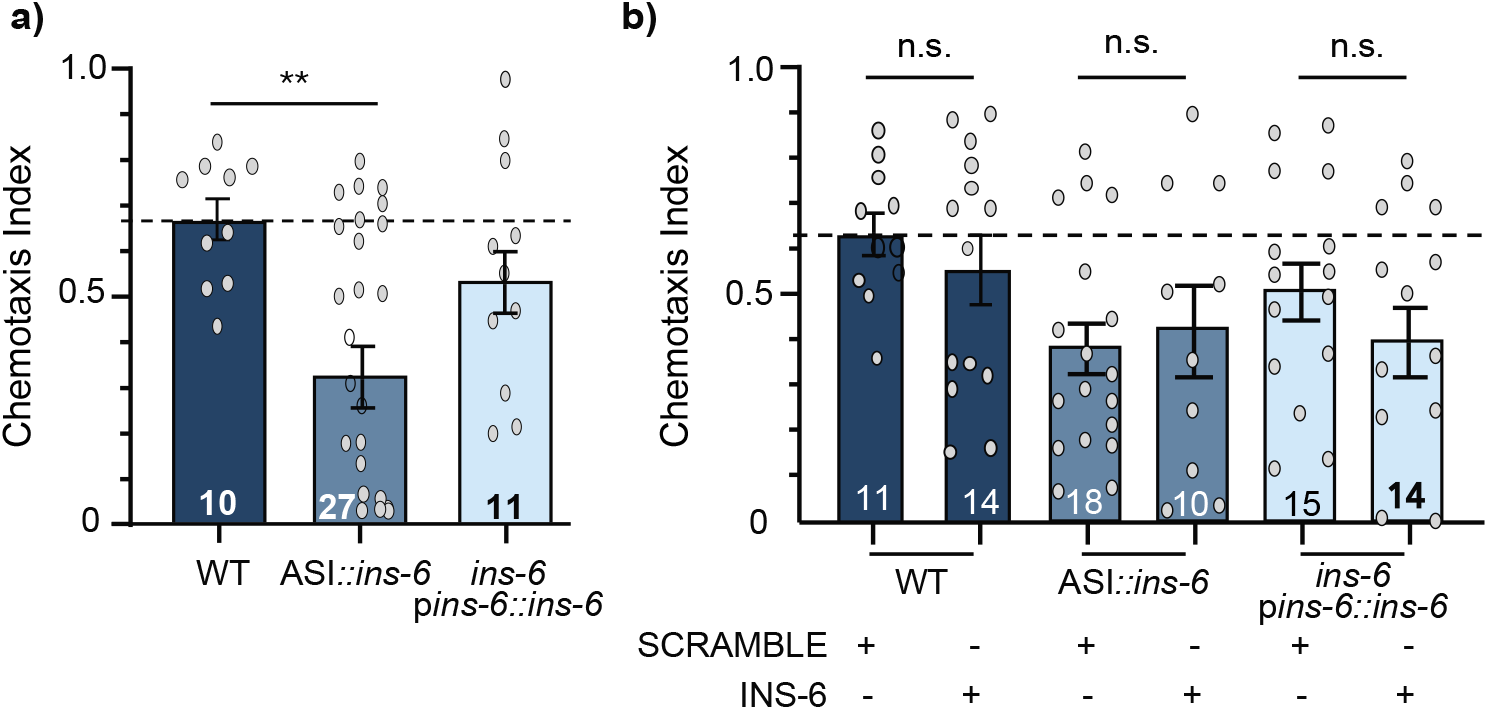

## Notes

### Competing Interest Statement

The authors have declared no competing interest.

