## Supplemental Information for "Non-Transgenic Functional Rescue of Neuropeptides"

a)

| Plasmid | Forward Sequence | Reverse Sequence |
|---|---|---|
| SCRAMBLE | GGGGACAAGTTTGTACAAAAAAGCAGGC TGGATGCGCTTTGGAAAACGTAATTCGA GCTCCACCGCGGTGGCGGCCGCTCTAGA ACTAGTGGATCCACCGGTTCCATGGCTA GCCACGCGCGTGGATCCCCCGGGCTGCA GGAAACGTAAATAACACCCAGCTTTCTT GTACAAAGTGGTCCCC | GGGGACCACTTTGTACAAGAAAGCTGG GTGTTATTTACGTTTCCTGCAGCCCGGG GGATCCACGCGCGTGGCTAGCCATGGAA CCGGTGGATCCACTAGTTCTAGAGCGGC CGCCACCGCGGTGGAGCTCGAATTACGT TTTCCAAAGCGCATCCAGCCTGCTTTTTT GTACAAACTTGTCCCC |
| TRH-1A | GGGGACAAGTTTGTACAAAAAAGCAGGC TGGATGCGCTTTGGAAAACGTAGAGGAC GAGAACTTTTCGGAAAACGTAAATAACA CCCAGCTTTCTTGTACAAAGTGGTCCCC | GGGGACCACTTTGTACAAGAAAGCTGG GTGTTATTTACGTTTTCCGAAAAGTTCTC GTCCTCTACGTTTTCCAAAGCGCATCCA GCCTGCTTTTTTGTACAAACTTGTCCCC |
| TRH-1B | GGGGACAAGTTTGTACAAAAAAGCAGGC TGGATGCGCTTTGGAAAACGTCGTGCCA ATGAACTTTTCGGTAAACGTAAATAACA CCCAGCTTTCTTGTACAAAGTGGTCCCC | GGGGACCACTTTGTACAAGAAAGCTGG GTGTTATTTACGTTTACCGAAAAGTTCA TTGGCACGACGTTTTCCAAAGCGCATCC AGCCTGCTTTTTTGTACAAACTTGTCCCC |
| INS-6 | CAATGCCACGAGCAAGTAGTGTTCCAG CACCAG | CTGGTGCTGGAACACTACTTGCTCGTGG CATTG |
| PDF-1A | GGGGACAAGTTTGTACAAAAAAGCAGGC TGGGTTCAGTTCGTAAAACGCAGCAACG CCGAGCTTATCAACGGACTCATCGGAAT GGATTTGGGAAAATTGTCAGCTGTCGGA AAACGCTGACACCCAGCTTTCTTGTACA AAGTGGTCCCC | GGGGACCACTTTGTACAAGAAAGCTGG GTGTCAGCGTTTTCCGACAGCTGACAAT TTTCCCAAATCCATTCCGATGAGTCCGTT GATAAGCTCGGCGTTGCTGCGTTTTACG AACTGAACCCAGCCTGCTTTTTTGTACA AACTTGTCCCC |
| PDF-1B | GGGGACAAGTTTGTACAAAAAAGCAGGC TGGGTTCAGTTCGTAAAACGCTCAAACG CGGAACTTATCAACGGTCTTCTCAGCAT GAACCTCAACAAATTGTCTGGAGCTGGT CGACGATGACACCCAGCTTTCTTGTACA AAGTGGTCCCC | GGGGACCACTTTGTACAAGAAAGCTGG GTGTCATCGTCGACCAGCTCCAGACAAT TTGTTGAGGTTCATGCTGAGAAGACCGT TGATAAGTTCCGCGTTTGAGCGTTTTAC GAACTGAACCCAGCCTGCTTTTTTGTAC AAACTTGTCCCC |

b)

| Peptide | Sequence (COOH-x-$NH_2$) |
|---|---|
| SCRAMBLE | NSKLHRGGGRSRTSGSTGSMASHARGSPGLQ |
| TRH-1A | GRELF |
| TRH-1B | ANELF |
| INS-6 | VPAPGETRACGRKLISLVMAVCGDLCNPQEGKDIATECCGNQCSDDYIRSACCP |
| PDF-1A | VQFVKRSNAELINGLIGMDLGKLSAVGKR |
| PDF-1B | VQFVKRSNAELINGLLSMNLNKLSGAGRR |

**Supplementary Table 1. Neuropeptide sequences used in this study. a)** Sequences of forward and reverse oligos used in generation of plasmids containing the peptide of interest for functional rescue of the neuropeptides **b)** Processed peptide amino acid sequences used for bacterial rescue experiments.

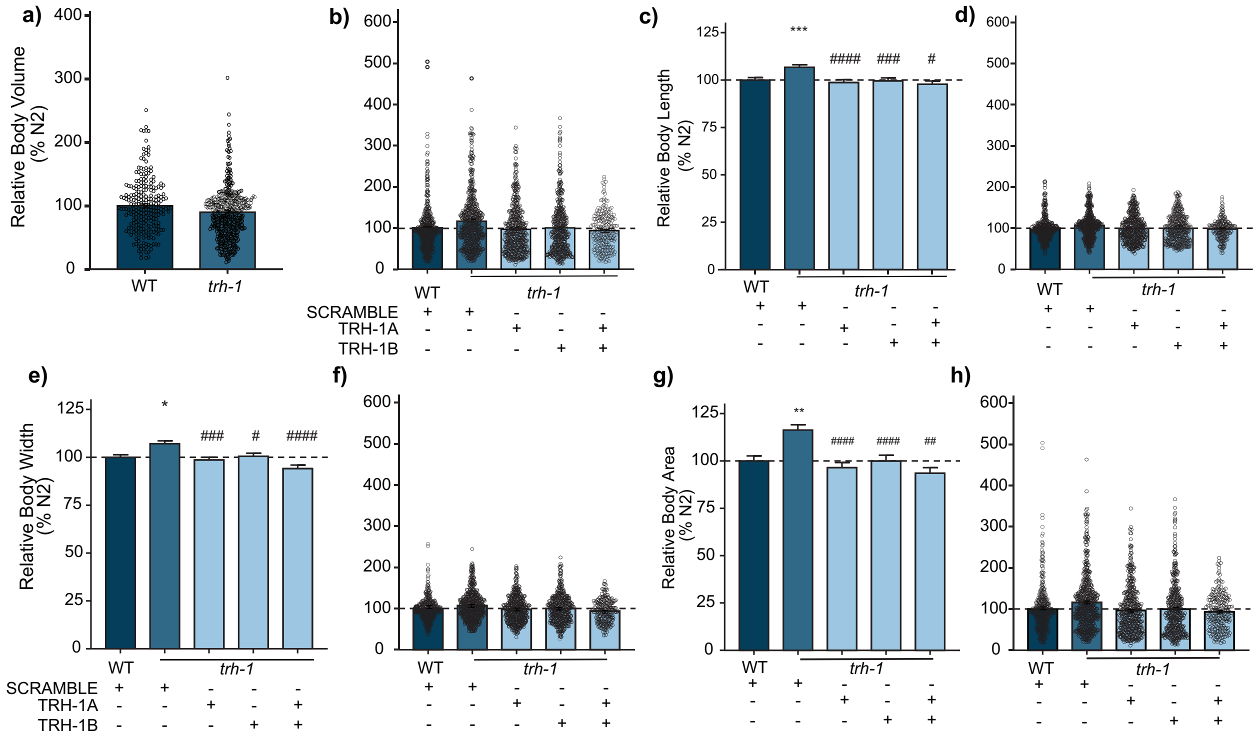


**Supplementary Figure 1. TRH-1 peptide feeding affects body length, width, and area of *trh-1* mutants. a)** Relative body volume of *E. coli* OP50 fed worms, displaying all data points (Mann-Whitney test, ** $p = 0.0012$). **b)** Relative body volume of peptide worms, all data points wild-type fed scramble, *trh-1* fed scramble, TRH-1A, TRH-1B, or both TRH-1A+B. **c)** Relative body length of peptide fed worms. **d)** Relative body length of peptide fed worms, all data points. **e)** Relative body width of worms fed peptide. **f)** Relative body width of peptide fed worms, all data points. **g)** Relative body area of peptide fed worms. **h)** Relative body area of peptide fed worms displaying all data points. *n* for all panels wild-type = 454, *trh-1* fed scramble = 559, fed TRH-1A = 483, fed TRH-1B = 441, fed TRH-1A+B = 221. Error bars denote SEM. (b-g) Kruskal-Wallis followed by Dunn's multiple comparison tests, */# $p < 0.05$, **/##, $p < 0.01$, ***/### $p < 0.001$, ****/#### $p < 0.0001$, *denote mutant worms compared to wild-type control, #denote mutant worm condition compared to mutant worms fed scramble peptide.

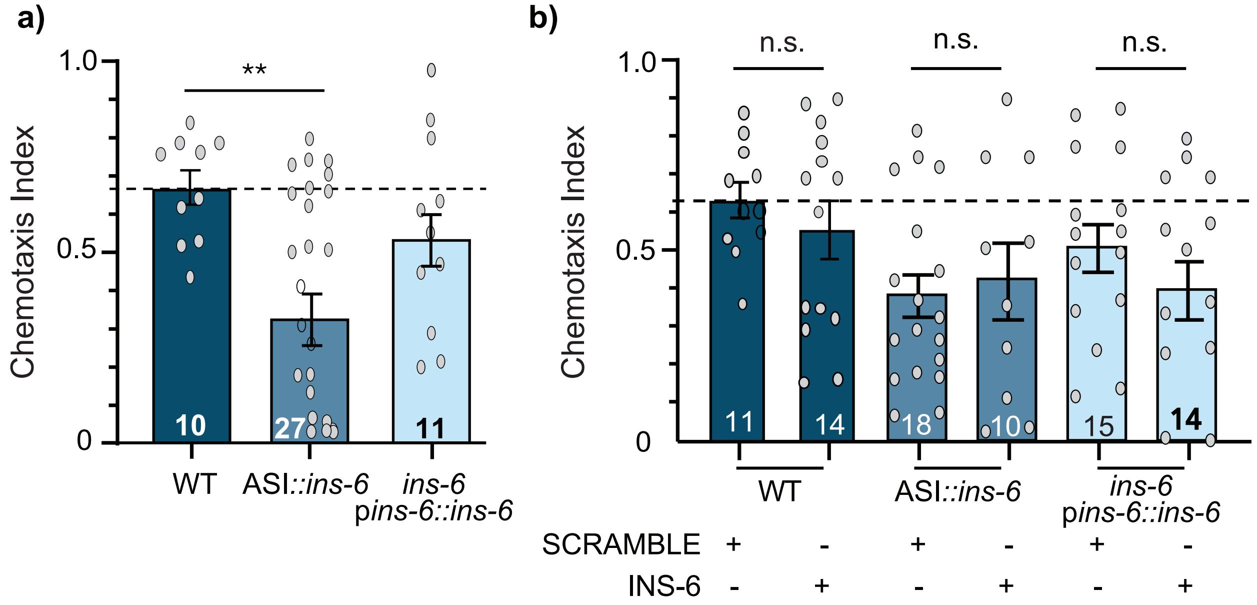


**Supplementary Figure 2. Chemotaxis Index of partially rescued *ins-6 lof* worms to 750mM NaCl without and with peptide feeding. a)** Chemotaxis Index to 750 mM NaCl of wild-type, *ins-6* rescued specifically in ASI neurons and a complete genetic rescue of *ins-6* animals. Rescue of ins-6 in ASI neurons does not result in NaCl chemotaxis, suggesting that this neuron does not play a role in this behavior. **b)** Chemotaxis Index of INS-6 peptide-fed wild-type and genetically rescued *ins-6* worms. Overexpression of INS-6 by feeding does not affect chemotaxis in wild type animals and the different rescue lines of *ins-6*. *n* values denoted in graphs. Error bars denote SEM. One-Way ANOVA, followed by Bonferroni's Correction. * $p < 0.05$, ** $p < 0.01$.

a)

| Genotype | Treatment | $P_L$ (confidence interval) | [*n*] |
|---|---|---|---|
| *him-5* | OP50 | 0.06553819 (0.05587592 , 0.07687131) | [72] |
| *him-5* | SCRAMBLE | 0.07727273 (0.06531852 , 0.09141473) | [55] |
| *him-5* | PDF-1A | 0.06979167 (0.05892111 , 0.08266776) | [60] |
| *him-5* | PDF-1B | 0.08712121 (0.0754002 , 0.1006643) | [66] |
| *him-5* | PDF-1A+B | 0.07720588 (0.06483647 , 0.09193512) | [51] |
| *pdf-1* | OP50 | 0.01785714 (0.0134571 , 0.02369585) | [84] |
| *pdf-1* | SCRAMBLE | 0.03537736 (0.02746855 , 0.04556329) | [53] |
| *pdf-1* | PDF-1A | 0.03286638 (0.02557212 , 0.04224128) | [58] |
| *pdf-1* | PDF-1B | 0.05821078 (0.04760713 , 0.07117621) | [61] |
| *pdf-1* | PDF-1A+B | 0.04473039 (0.03556126 , 0.0562637) | [51] |

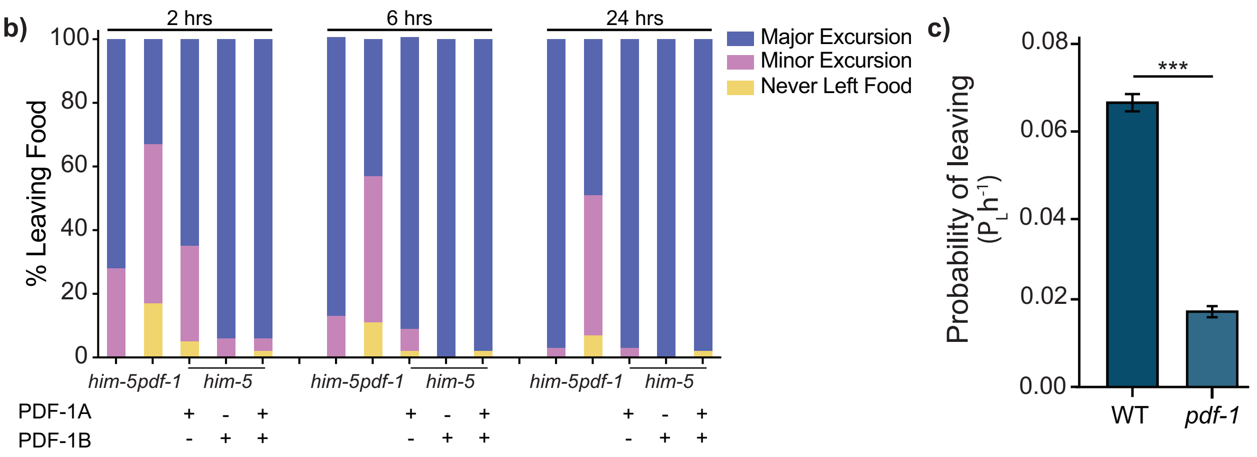


**Supplementary Figure 3. Leaving behavior of *him-5* and *pdf-1* males fed PDF-1 peptides. a)** Table displaying the calculated Probability of Leaving $P_L$ for each peptide feeding condition in *him-5* and *pdf-1* males. The table also depicts the confidence interval of the data and the sample size for each condition. **b)** Percentage of *him*-5 males leaving food after being fed PDF-1 peptides on *E. coli* OP50. PDF-1B feeding results in complete rescue of food-leaving behavior of *pdf-1 lof males* at all three timepoints **c)** Computation of Probability of leaving for *him-5* and *pdf-1* males fed *E. coli* OP50, *n* noted in **a) [*n*]** column. Error bars denote SEM. **b)** ANOVA, followed by Bonferroni's Correction, **c)** Two-tailed *t*-test of samples of equal variance. *** $p < 0.001$.
